# The asgardarchaeal-unique contribution to protein families of the eukaryotic common ancestor was 0.3%

**DOI:** 10.1101/2021.02.09.430432

**Authors:** Michael Knopp, Simon Stockhorst, Mark van der Giezen, Sriram G. Garg, Sven B. Gould

## Abstract

Ever since the first report of a new archaeal lineage, the asgardarchaea, their metagenome analyses have encouraged continued speculations on a type of cell biology ranging between that of prokaryotes and eukaryotes. While it appears a tempting notion, recent microscopic images of an asgardarchaeon suggest otherwise. We inspected the origin of eukaryotic protein families with respect to their distribution across bacteria and archaea. This reveals that the protein families shared exclusively between asgardarchaea and eukaryotes amounts to only 0.3% of the protein families conserved across all eukaryotes. Asgardarchaeal diversity is likely unrivaled across archaea, but their cell biology remains prokaryotic in nature and lends support for the importance of endosymbiosis in evolving eukaryotic traits.

**Summary:** The difference between pro- and eukaryotic biology is evident in their genomes, cell biology, and evolution of complex and macroscopic body plans. The lack of intermediates between the two types of cells places the endosymbiotic acquisition of the mitochondrion through an archaeal host at the event horizon of eukaryote origin. The identification of eukaryote specific proteins in a new archaeal phylum, the asgardarchaea, has fueled speculations about their cellular complexity, suggesting they could be eukaryote-like. Here we analyzed the coding capacity of 150 eukaryotes, 1000 bacteria, and 226 archaea, including the only cultured member of the asgardarchaea, Candidatus *Prometheoarchaeon syntrophicum* MK-D1. Established clustering methods that recover endosymbiotic contributions to eukaryotic genomes, recover an asgardarchaeal-unique contribution of a mere 0.3% to protein families present in the last eukaryotic common ancestor, while simultaneously suggesting that asgardarchaeal diversity rivals that of all other archaea combined. Furthermore, we show that the number of homologs shared exclusively between asgardarchaea and eukaryotes is only 27 on average. Genomic and in particular cellular complexity remains a eukaryote-specific feature and, we conclude, is best understood as the archaeal host’s solution to housing an endosymbiont and not as a preparation for obtaining one.

## Introduction

Four billion years of prokaryotic evolution has only once resulted in the emergence of highly compartmentalized cells and eventually macroscopic body plans: following the origin of eukaryotes through endosymbiosis. The analysis of core eukaryotic features such as the nucleus, mitochondria, sex and meiosis, compartmentalization and dynamic membrane trafficking, and virtually all of the associated protein families, consistently point to their presence in the last <u>e</u>ukaryotic <u>c</u>ommon <u>a</u>ncestor (LECA) (Fritz-Laylin et al. 2010, Koumandou et al. 2013, Koonin et al. 2013, Garg and Martin 2016). We possess a reasonable understanding of the basic cellular features and coding capacity of LECA, owing to the growing number of genome sequences spanning all of eukaryotic diversity. All eukaryotes stem from a single ancestor that in terms of cellular and genomic complexity rivaled those of extant eukaryotic supergroups (Fritz-Laylin et al. 2010, Koumandou et al. 2013, Koonin et al. 2013). It is certain that LECA was a product of the integration of an alphaproteobacterium into an archaeal host following endosymbiosis (Lane 2011, Blackstone 2013, Martin et al. 2015, Dacks et al. 2016, Spang et al. 2019).

Through the description of the asgardarchaea, current debates once again concern the cellular complexity of the host that came to house the endosymbiont and what contribution the mitochondrion could have played in establishing the eukaryotic cell (Dacks et al. 2016, Martin et al. 2017). Asgardarchaea, a novel phylum assembled from metagenome data, are viewed as bridging the gap between pro- and eukaryotic cells, because they encode proteins homologous to eukaryotic ones that are e.g. involved in intracellular vesicle trafficking and the regulation of actin cytoskeleton dynamics (Spang et al. 2015, Zaremba-Niedzwiedzka et al. 2017, Neveu et al. 2020). The cellular complexity of the host cell that acquired the alphaproteobacterial endosymbiont has been a matter of speculation ever since the realization that endosymbiosis was pivotal in the transition to eukaryotic life. Modern models of eukaryogenesis differ regarding the timing of mitochondrial acquisition, the extent of the cellular complexity of the host, and the selective reasons provided for explaining the presence, function, and emergence of eukaryotic traits prior or ensuing endosymbiosis (O’Malley 2010, Martin et al. 2015, Gould et al. 2016, Tria et al. 2019, Vosseberg et al. 2020).

Understanding the steps of eukaryogenesis is a demanding intellectual challenge that explores the past of life and one of its most radical transitions. It holds the key to understanding the steps towards cellular complexity, the timing of mitochondrial entry, and what limits prokaryotes to frequently evolve eukaryote-like complexity. Was eukaryogenesis really a matter of luck (Booth and Doolittle 2015) and how important was the energetic superiority provided by the mitochondrion to the host cell (Lane and Martin 2010, Lynch and Marinov 2015, Lane and Martin 2016)? Any model that views endosymbiosis as some kind of terminal coincidence on the evolutionary roadmap to the eukaryotic domain of life needs to explain the singularity that is eukaryogenesis and the lack of comparable complexity among prokaryotes.

A consistent motivation for speculating on the archaeal host cell’s grade of complexity is trying to understand whether the host cell was phagocytotic or not (Cavalier-Smith 1987, Yutin et al. 2009, Martijn and Ettema 2013, Martin et al. 2017). This is complicated by the description of a phagocytosis-like process in a planctomycete (Shiratori et al. 2019) and the conflicting evidence for intracellular prokaryotic endosymbionts in the absence of phagocytosis (Fenchel and Bernard 1993, Emblay and Finlay 1993, Schmid 2003, Zientz et al. 2004, Duplessis et al. 2004, Thacker 2005, Woyke et al. 2006, Husnik et al. 2013). It has also been concluded that asgardarchaea are not phagocytotic (Burns et al. 2018), although they encode actin-regulating profilins (Akil and Robinson 2018), small Rab-like GTPases (Surkont and Pereira-Leal 2016), and prototypic SNARE proteins (Neveu et al. 2020). Phagocytosis might have evolved multiple times independently (Yutin et al. 2009, Mills 2020) and is a mode of feeding, which is incompatible with the syntrophic foundation that underpins eukaryogenesis (Martin et al. 2015, Spang et al. 2019, Martin and Müller 1998, Vellai et al. 1998, Imachi et al. 2020). A sole focus on this single eukaryotic trait might distract and furthermore discounts the complexity of the transition that was involved. What is certain is that images of an asgardarchaeon, Candidatus *Prometheoarchaeum syntrophicum* MK-D1, reveal cells with typical archaeal morphology, half a micron in diameter, with obligate syntrophy, and devoid of intracellular complexity (Imachi et al. 2020).

Here we clustered the available genomes of 150 eukaryotes, 1000 bacteria and 226 archaea (including asgardarchaea metagenomic assemblies, and for comparison the complete genome of the cultured Candidatus *P. syntrophicum* strain MK-D1) in order to evaluate the asgardarchaeal-unique contribution to eukaryogenesis that is understood as support for early cellular complexity in asgardarchaea.

## Results

In order to evaluate to what degree asgardarchaea bridge the prokaryotic and eukaryotic protein families, we performed a global comparison of clustered gene families across eleven asgardarchaeal metagenome-assembled genomes (MAGs), the closed MAG of the cultured asgardarchaeon Candidatus *P. syntrophicum* MK-D1, 214 other archaea, 1,000 bacteria, and 150 eukaryotes. Protein families for 150 eukaryotes were taken from Brueckner and Martin (Brueckner and Martin 2020), which included 239,012 clusters. We further clustered proteins from the prokaryotes, resulting in 352,384 bacterial clusters and 49,855 archaeal clusters. Subsequently, the eukaryote and prokaryote clusters were merged in a reciprocal best cluster approach previously described in Ku et al. 2015 (Ku et al. 2015), yielding Eukaryote-Prokaryote-Clusters (EPCs). These EPCs contained proteins from eukaryotes and proteins from either archaeal (Eukaryote-Archaea clusters, EA) or bacterial (Eukaryote-Bacteria clusters, EB), or both (Eukaryote-Archaea-Bacteria clusters, EAB). This approach yielded 2,590 EPCs, of which 867 or 33.5% (330 EA clusters + 537 EAB clusters; Suppl. Table 1) contained an archaeal component. Among these 867 EPCs, asgardarchaeal protein sequences, including those of Candidatus *P. syntrophicum*, were present in about 75% of the protein families (75% of EAB clusters and 75.2% of EA clusters).

A presence-absence pattern (PAP) of all 867 protein families with archaeal contribution to the EPCs, including the 537 protein families present in all domains is shown in Fig. 1 (and Suppl. Table 1). While gene distributions among eury- and crenarchaeota is highly similar, those of the asgardarchaea are patchier and more diverse. Among all of our 239,012 eukaryotic clusters, we could identify only six EA clusters with asgardarchaeal-unique contributions to eukaryotes (Suppl. Table 2), representing 0.0025% of all extant eukaryotic diversity. To calculate the asgardarchaeal-unique contribution to LECA, we filtered the eukaryotic clusters for those that include at least one representative of each of the six eukaryotic supergroups, resulting in 1880 LECA clusters and consequently an asgardarchaeal-unique contribution of 0.3191%.

**Fig 1:**
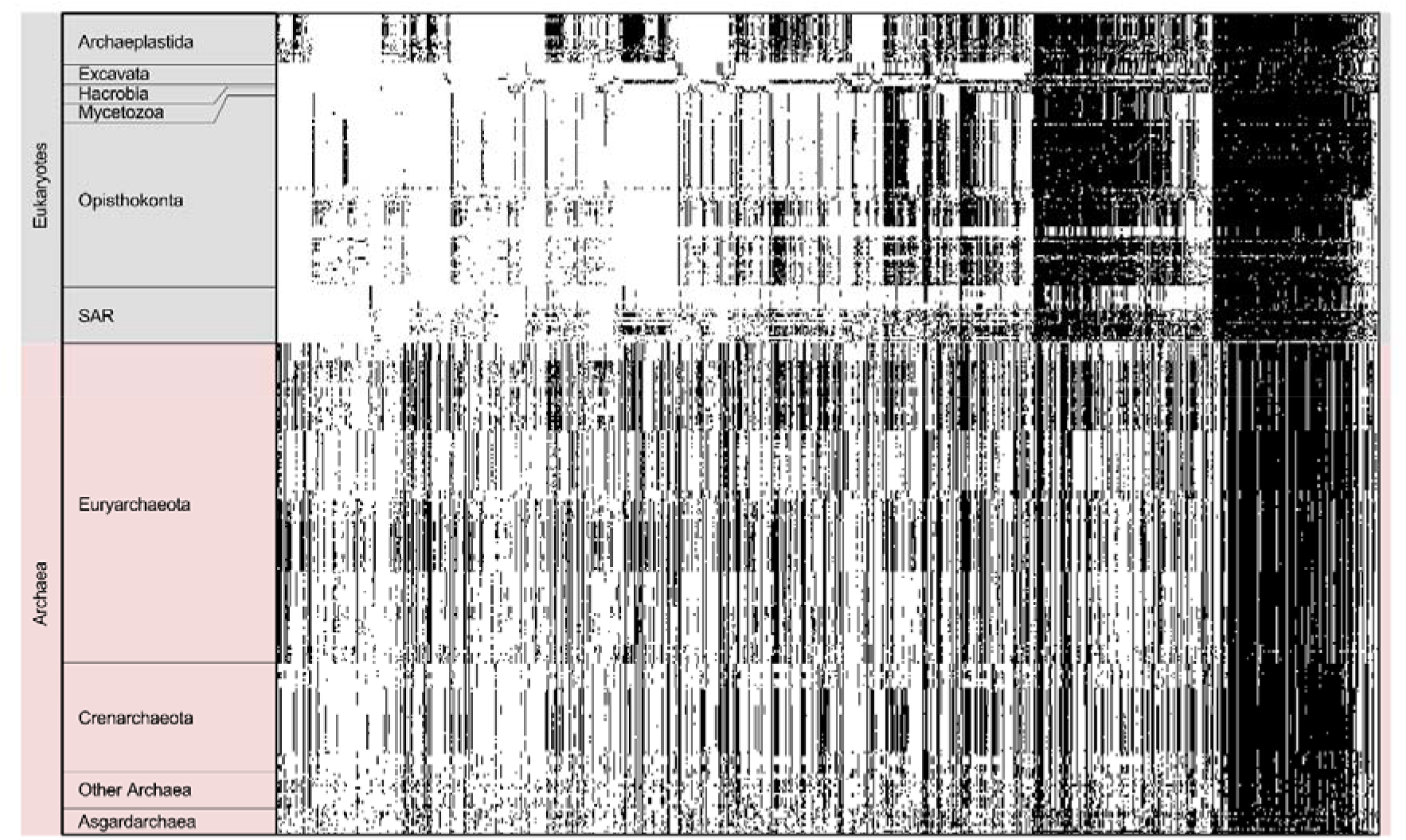
Presence-absence pattern (PAP) of all 867 eukaryote-prokaryote clusters (EPCs) with archaeal contribution among the investigated 150 eukaryotes, 212 archaea and twelve asgardarchaea. The protein families were sorted by their distribution among six eukaryotic supergroups (SAR; Stramenopila, Alveolata, Rhizaria), their presence is indicated in black along the X-axis. The group of “Other Archaea” is comprised of all archaea within our database that have less than 15 members. The distribution of these EPCs among the asgardarchaea does not reveal a particular pattern such as that seen for the other archaeal groups, demonstrating their large genetic differences.

But how closely are the asgardarchaea related to each other with respect to other archaea? To investigate their kinship, we constructed protein families of only the asgardarchaeal protein sequences, which generated a set of 5,837 protein families. The distribution of these protein families among the asgardarchaea reveals a pronounced diversity, with only a small proportion of the protein families being shared across all of them, which suggests one is dealing with a kind of superphylum (Fig. 2a). This is evident from a comparison wherein, we clustered all protein sequences of 14 members of the genus Methanococci and 52 crenarchaeote members from the TACK superphylum (Fig. 2b and Fig. 2c), revealing 2,592 and 12,871 protein families, respectively.

**Fig 2:**
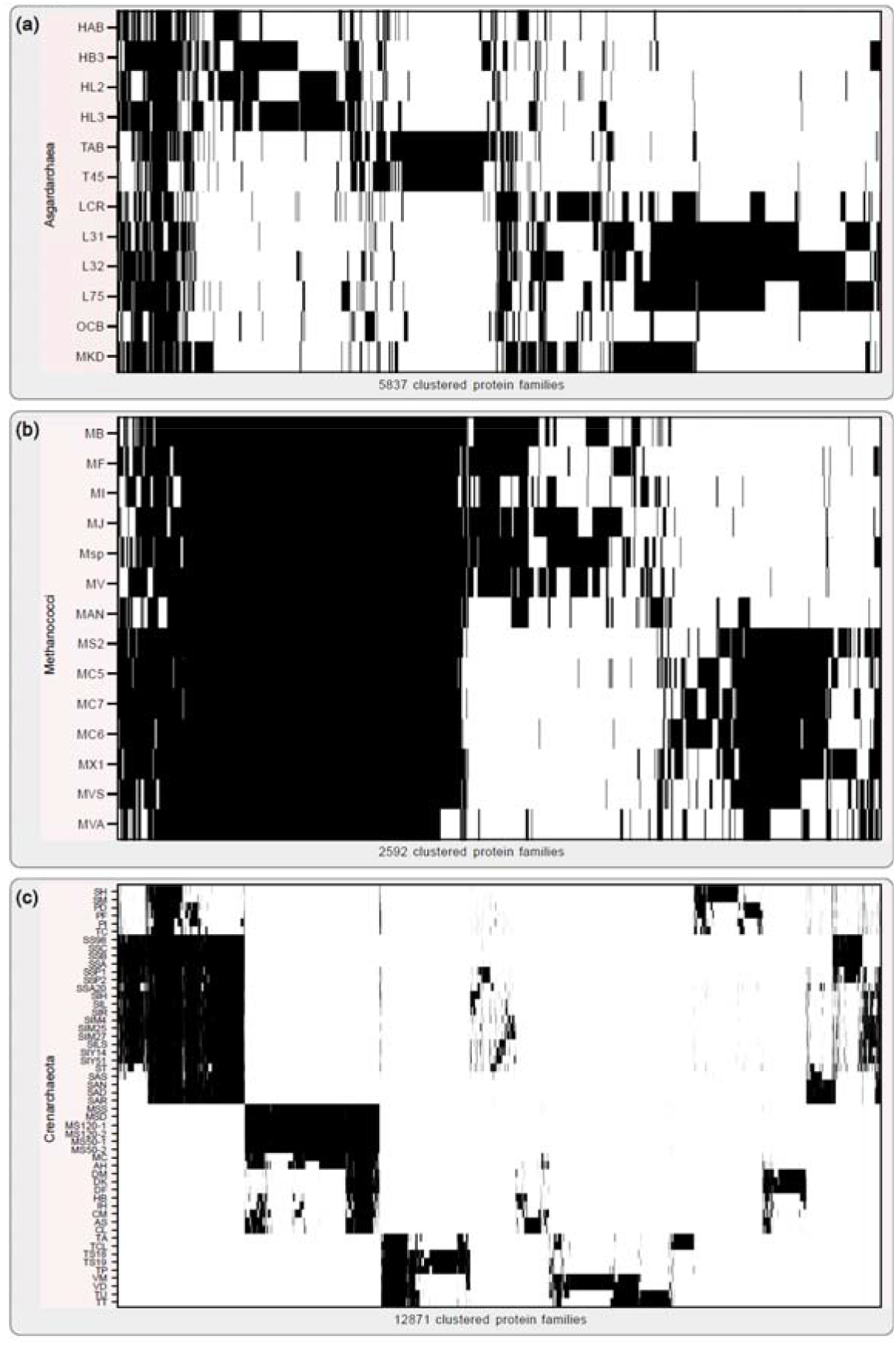
Archaeal protein family distributions. **(a)** Distribution of all 5,837 calculated asgardarchaeal protein families among the investigated asgardarchaea. The protein families were obtained by globally comparing all asgardarchaeal protein sequences in a pairwise all-vs-all Diamond BLASTp approach including subsequent clustering via MCL. The result was sorted via hierarchical clustering along the X-axis. The vast majority of protein families is not shared among all asgardarchaea but they are rather specific to the individual taxon. Candidatus Prometheoarchaeon syntrophicum MK-D1 shares the highest number of its protein families with members of the Lokiarchaeota. **(b)** For comparison, we calculated protein families for 14 members of the class Methanococci in the same manner, generating a total of 2,592 protein families. Hierarchical clustering revealed a striking difference in protein families shared between members within the two clusterings, underlining the investigated asgardarchaea’s diversity among each other. **(c)** On the contrary, clustering of 52 members of the crenarchaeota, a highly diverse taxonomic group, results in 12,871 protein families. The matrix was sorted along the X- and Y-axis via hierarchical clustering. All Abbreviations used are listed in Suppl. Table 6.

A key difference between archaea and eukaryotes is the difference in the number of protein families. In a previous study, 239,012 protein families were generated on the basis of all pairwise protein sequence comparisons of 150 eukaryotic proteomes (Brueckner and Martin 2020). The same approach yields a total of 49,855 protein families on the basis of all pairwise protein sequence comparisons between 226 archaea, including 11 asgardarchaea and Candidatus *P. syntrophicum* MK-D1. The patchy distribution of the asgardarchaeal EPCs (Fig. 1) and the high diversity between each other (Fig. 2a) prompts a closer look, but the vast difference in the number of protein families reflects the sudden inflation of protein family emergence at the origin of eukaryotes.

Since many asgardarchaeal protein sequences could not be clustered due to a lack of homology, we conducted an analysis of BLASTp hits against a bacterial database consisting of 5,443 proteomes, an archaeal database consisting of 212 proteomes (excluding asgardarchaea) and a eukaryotic database consisting of 150 proteomes. For all investigated asgardarchaea and a selection of twelve well known and diverse archaea and bacteria each, as well as eight eukaryotes, we quantified the amount of sequences with homologs in one of these databases, any combination of these, or no significant homologs at all (Fig. 3a), while ignoring hits from the same genus to counter database composition biases. For asgardarchaea, only 27 protein sequences on average have homologs unique to eukaryotes (Fig 3a, magnified area), supporting the initial EPC analysis which only uncovered six protein families that were unique to eukaryotes and asgardarchaea. Furthermore, this test reveals that high proportions of the asgardarchaeal proteomes do not retrieve significant hits in our three tested databases (Fig. 3a, grey bars). In a few cases, such as for example for Candidatus *Heimdallarchaeota LC_3* or Candidatus *Lokiarchaeota archaeon CR_4*, the number of proteins for which no homology was detected in any other species, exceeds half of the respective genome’s coding capacity (Fig. 3a).

**Fig 3:**
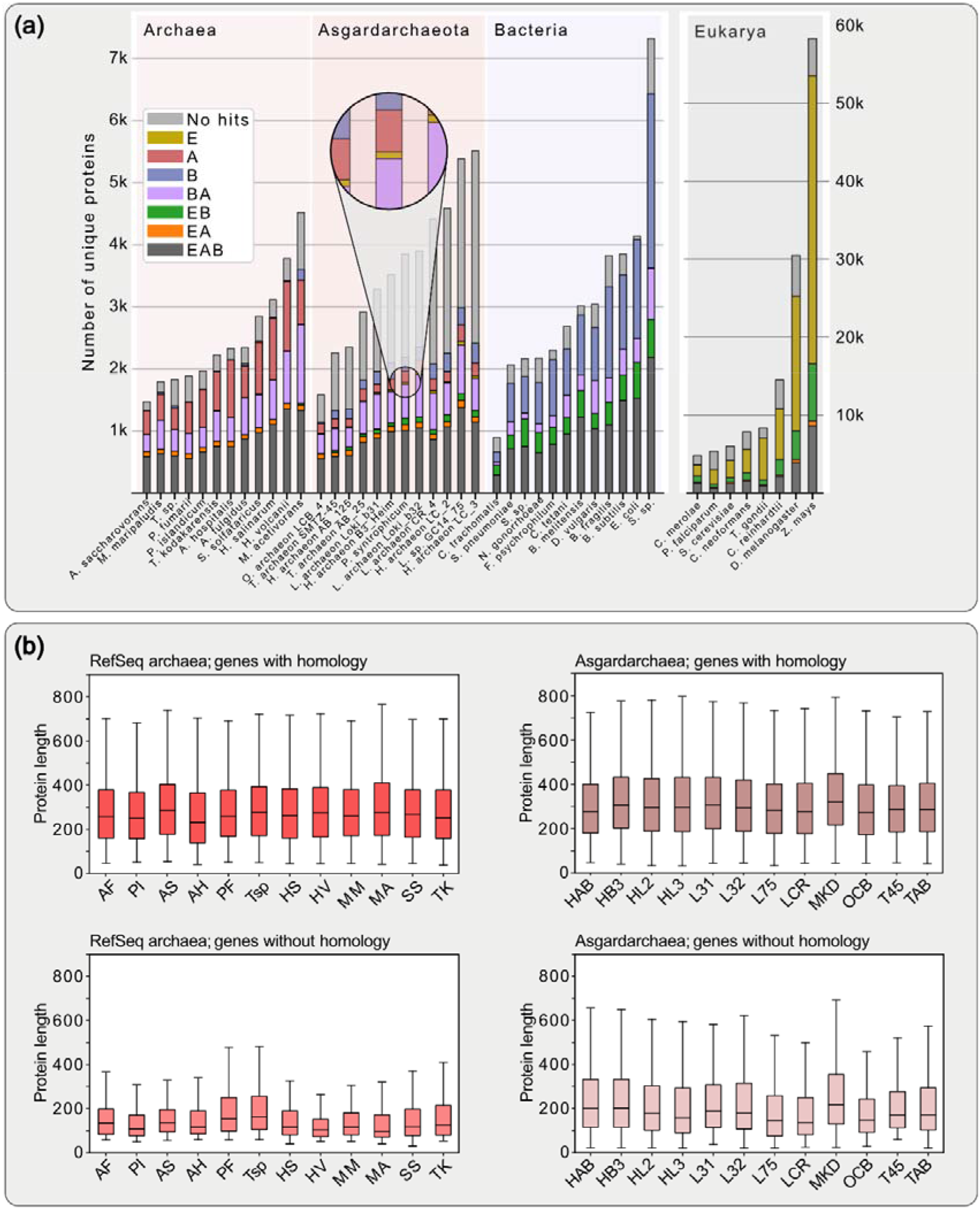
Distribution of protein homologs and length comparison of archaeal genes with and without homology. **(a)** BLAST hit analysis comparing the proteomes of known archaea and bacteria as well as the twelve investigated asgardarchaea. Sequences were blasted against a prokaryotic database comprising 5,655 prokaryotic genomes (including the 1000 bacteria and 212 archaea which were used for protein family calculation) and a eukaryotic database comprised of 150 genomes. For each organism we quantified the amount of sequences with subjects among eukaryotes, bacteria or archaea only, or any combination of these three. To counter possible biases in database composition, hits from the same genus were excluded. E, Eukaryotes; A, Archaea; B, Bacteria; AB, Archaea-Bacteria; et cetera. **(b)** For proteins with- or without database hits, protein length distributions are shown as box and whiskers plots. This reveals a noticeable difference in protein lengths of sequences without database hits from only the asgardarchaea compared to their sequences with database hits, and in comparison, to those of Refseq archaea. p-values of FDR-corrected, pairwise double-sided Kolmogorov-Smirnov tests in Suppl. Table 7. All used abbreviations are listed in Suppl. Table 6.

We plotted the protein sequence length distributions for each proteome, separating hits with and hits without significant database hits (Fig. 3b). While the length distributions of sequences with significant database hits were comparable between the asgard- and the RefSeq archaea, the distributions of hits without significant database hits show a major difference. (Fig. 3b). Furthermore, simulating missing data by ignoring not only hits from the same genus but also the same phylum produced a more similar result for the reference organisms and asgardarchaea (Suppl. Fig.1), recapitulating the extensive diversity within the asgardarchaea.

## Discussion

The identification of asgardarchaea from deep-sediment metagenome data (Seitz et al. 2016; Zaremba-Niedzwiedzka et al. 2017) provides valuable new information from which to re-evaluate key issues surrounding the tree of life and the emergence of its eukaryotic branch. To begin with, the iconic tree that introduced three aboriginal lineages (Woese and Fox 1977) might require a revision. Phylogenomic analysis of asgardarchaea provides evidence for a two-domains tree and the emergence of the host cell lineage of eukaryogenesis from within the archaeal domain (Cox et al. 2008, McInerney et al. 2014, Hug et al. 2016, Williams et al. 2020) with some skepticism, however, remaining (Forterre 2015, Da Cunha et al. 2018, Liu et al. 2020). Parallel to the discovery of the asgardarchaea were immediate speculations regarding their cellular complexity (Dacks et al. 2016, Pittis and Gabaldón 2016, Rout and Field 2017, Akıl and Robinson 2018, Zachar et al. 2018, Neveu et al. 2020) and a faith of having identified the missing link between pro- and eukaryotic biology based on the identification of a few <u>e</u>ukaryote <u>s</u>ignature <u>p</u>roteins (ESPs), that we find to be only 27 on average. In light of these numbers, the potential of these archaea to display eukaryote-like cell complexity is hard to maintain.

Our analysis confirms a patchy distribution of ESPs among asgardarchaea (Dacks et al. 2016, Zaremba-Niedzwiedzka et al. 2017, Imachi et al. 2020, Liu et al. 2020, Klinger et al. 2016, Inoue et al. 2020, Bulzu et al. 2019) (Fig. 1). While of course the archaeal host brought in 1000s of genes, the unique contribution of this new archaeal lineage to the eukaryotic protein families is substantially less than what one might infer from the original metagenome reports and subsequent interpretations (Dacks et al. 2016, Pittis and Gabaldón 2016, Rout and Field 2017, Zachar et al. 2018, Akıl and Robinson 2018, Neveu et al. 2020, Vosseberg et al. 2020). The irregular gene distribution (Fig. 2a) might reflect differential gene loss upon the segregation of the common ancestor of asgardarchaea and the archaeal host cell lineage (Eme et al. 2017). Considering the role of pangenomes in the transformation of prokaryotic lineages and the conquering of ecological niches (McInerney et al. 2017), pangenomes offer a complementary explanation to the differential loss of genes. The archaeal ancestor of eukaryotes might have tapped a shared gene pool more extensively than the sister lineages leading to extant asgardarchaea.

The notion that asgardarchaeal contributions to eukaryotes were higher only to be eventually replaced by bacterial (endosymbiotic) contributions might be brought up (Pittis and Gabaldon 2016; Eme et al. 2017; Vosseberg et al. 2020). The absence of extant archaeal relatives with similarly higher ESPs, however, indicates that archaea neither have the necessity nor the selective pressure for maintaining ESPs in the absence of an endosymbiont. Any theory that hinges upon a larger presence of ESPs in archaea or bacteria prior to mitochondrial endosymbiosis ignores the complete lack of “accumulated ESPs” in extant prokaryotes to a degree that even remotely matches that of any given eukaryotic lineage. Extinction and absence of geological conditions that promoted eukaryote origins considered for ESPs, must also be considered for any genes that are currently thought to be recent independent gene transfer events from prokaryotes to eukaryotes and not of endosymbiotic (mitochondrial) origin.

Our protein family clustering method, which readily detects the mitochondrial contribution (and the cyanobacterial contributions in the case of the Archaeplastida; Suppl. Fig. 2) failed to detect a comparable asgardarchaeal-unique contribution. A stacked bar diagram puts gene family cluster contribution in each lineage into a global perspective (Fig. 3a; Supp. Fig 1). There is a small proportion of eukaryotic homologs (E, mustard yellow) visible, e.g. in the Candidatus *P. syntrophicum* MK-D1, but it is substantially smaller in comparison to the eukaryote-bacteria (EB, green) specific homologs evident in eukaryotes (and bacteria *vice versa*) that reflects the mitochondrial contribution to eukaryogenesis (Brueckner and Martin 2020). Neglecting the surprisingly low number of asgardarchaeal-unique homologs to eukaryotic genomes, our analysis demonstrates that asgardarchaea are among the most genetically diverse group of archaea when comparing it to the genus Methanococci and the phylum Crenarchaeota (Fig. 2). The odd length distributions of the asgardarchaeal proteins with no homology (Fig. 3b) are strange as well, since protein length across pro- and eukaryotes is usually well conserved (Xu et al. 2006). This could hint at an assembly and/or binning issue, which was also observed regarding the anomalous phylogenetic behavior of their ribosomal proteins and concatenated gene trees (Da Cunha et al. 2018; Garg et al. 2021). If not, it is a biological phenomenon absent in other sequenced prokaryotes.

Considering the amount of data gathered in just the few years (Imachi et al. 2020, Liu et al. 2020, Klinger et al. 2016, Inoue et al. 2020, Bulzu et al. 2019, Villanueva et al. 2016), it is surprising asgardarchaea have escaped identification for so long. Their habitats had been sampled before, so it is likely that the method used and maybe an obligate dependency on syntrophy, hindered culturing except for one imposing exception (Imachi et al. 2020). Dedicated phylogenomic efforts are necessary to resolve their taxonomic classification, while only culturing can picture their cell morphology.

Analyses of asgardarchaeal ESPs shows they have the potential to function similar to their eukaryotic homologs in a eukaryotic system (Neveu et al. 2020, Akıl and Robinson 2018, Rout and Field 2017, Klinger et al. 2016), but cross-kingdom inferences have their limits (Dey et al. 2016). The analysis of archaeal small GTPases (Surkont and Pereira-Leal 2016) and homologs of ESCRT proteins, the CDVs (Lindås et al. 2008, Lindås and Bernander 2013, Caspi and Dekker 2018), serve as examples. One needs to interpret asgardarchaeal ESPs in their prokaryotic context and in cells lacking an endosymbiont. The first images of an asgardarchaeon, those of Candidatus *P. syntrophicum* MK-D1, and its dependency on a bacterial partner (Imachi et al .2020) define the current standard from which to plot eukaryogenesis and the steps leading to eukaryotic cell- and genome complexity.

The identification of the asgardarchaea and the culturing of one representative represent an important milestone in micro- and evolutionary biology. Their phylogenetic analysis echoes two previously predicted outcomes: (i) eukaryotes to branch from within archaea, solidifying the two-domains tree of life, and (ii) that the closer we zoom in on the two prokaryotic partners from which eukaryotes evolved, the higher the number of otherwise eukaryote-typical genes we identify in prokaryotes. The description of Candidatus *P. syntrophicum* MK-D1 (Imachi et al. 2020) reminds us to not conflate genotypic with phenotypic complexity. This predicts that future asgardarchaea we see cultured will lack eukaryotic traits, too, and most, if not all, will depend on syntrophy. Our analysis also predicts that much of asgardarchaeal diversity remains to be described, but that the gap between the pro- and eukaryotic protein families will remain decisive and to change little. Placing the endosymbiotic event and the energetic benefit offered by mitochondria to fuel the transition early in eukaryogenesis, explains the lack of physical evidence for eukaryote-like complexity in asgardarchaea despite them encoding ESPs. It offers a comprehensive full-service theory for the singularity that is the origin of the eukaryotic cell that mitochondria-late models fail to provide.

## Acknowledgements

SBG would like to thank the German research council (267205415 – SFB 1208) and the VolkswagenStiftung (Life) for funding. SGG was furthermore supported by the Moore–Simons Project on the Origin of the Eukaryotic Cell GBMF9743. Computational infrastructure and support were provided by the Centre for Information and Media Technology at HHU Düsseldorf.

## Author Contributions

SGG, MK and SBG conceived the analysis, which was carried out predominantly by MK but also SS. SGG, MK, MvdG and SBG wrote the manuscript, whose final version was approved all authors.

## Methods

### Calculation of protein families

Prokaryotic gene families were calculated from complete genomes of 1000 bacteria and 226 archaea of the Refseq database (Version September 2016) including eleven representatives of the asgardarchaea^13^ and Candidatus *P. syntrophicum* MK-D1 (Imachi et al. 2020), separately. Bacterial and archaeal protein families were calculated via MCL (van Dongen 2000, Enright 2002) (--abc -scheme 7) from all reciprocal best BLAST hits with pairwise global identities (Rice et al. 2000) of at least 25% identity and a maximum e-value of 1×10^−10^ among all investigated bacteria and archaea, respectively. Eukaryotic gene families on the basis of 150 eukaryotic genomes were calculated as part of a previous study (Brueckner et al. 2020). Prokaryotic and eukaryotic cluster were combined into EPCs if at least 50% of all sequences of a eukaryotic cluster had their best hit in a prokaryotic cluster and vice versa according to the ‘reciprocal best cluster approach’ described in Ku et al. 2015 (Ku et al. 2015). All proteomes used in this study are listed in Supplementary Table 5. Protein families of Methanococci and Crenarchaeota were calculated from all pairwise global identities of all protein sequences, using the above-mentioned identity and e-value cutoffs, via MCL (van Dongen 2000, Enright 2002).

### BLAST hit analysis

We performed a Diamond BLASTp (Buchfink et al. 2015) hit analysis on all protein sequences of the eleven asgardarchaeal proteomes including MK-D1. We compared the results to twelve bacterial and archaeal proteomes each, plus eight eukaryotic ones. We blasted all protein sequences of the chosen proteomes against our database of 5655 prokaryotic and 150 eukaryotic proteomes. For each proteome, we quantified the number of sequences that showed significant hits (at least 25% identity and a minimal e-value of 1×10^−5^) within bacteria, archaea, eukaryota or any combination of these three. To counter overrepresentation of some genera within our database, we excluded hits of the same genus for all tested protein sequences.

### InterProScan ESP analysis

InterProScan version 5.39-77.0 (Quevillon et al. 2005) with standard parameters was used to annotate all asgardarchaeal proteomes, the MK-D1 proteome and all 212 archaeal proteomes within our prokaryotic database. As in Zaremba et al. 2017 (Zaremba-Niedzwiedzka et al. 2017), we searched for InterPro-Identifiers that correspond with Eukaryote-specific-proteins and plotted the results of all investigated asgard archaea together with the results of 14 model Refseq archaea for comparison. (see Suppl. Figure 3)

### LECA cluster filtering

Since eukaryotic inheritance is strictly vertical the eukaryotic protein families were filtered for families that contained at least one protein sequence from one member of each of the six supergroups included in our dataset of 150 eukaryotes, being *Archaeplastida, Opisthokonta, SAR, Hacrobia, Excavata* and *Mycetozoa* resulting in 1880 protein families passing this criterion.

Suppl. material titles and legends

**Suppl. Fig. 1**: **BLAST hit analysis**. The BLAST hit analysis underlying Fig. 3a was redone, this time ignoring all hits from the same phylum. This was done in order to test for the asgardarchaeas’ taxonomic rank, since the amount of protein sequences without any significant database hits, could either stem from poor sequence quality or just from the fact that asgardarchaea are inherently different from all archaea we know so far. As expected, the amount of Refseq archaeal protein sequences without significant database hits did increase but the effect was not sufficient to recover the same proportion of “unknown” protein sequences as we see in asgardarchaea.

**Suppl. Fig. 2: PAPs of sets of Eukaryote-Prokaryote-Clusters. (a)** Presence-Absence pattern of all 537 EPCs containing members from eukaryotes, bacteria and archaea. **(b)** Presence-Absence pattern of all 1723 EPCs containing only proteins from eukaryotes and bacteria. The cyanobacterial contribution to the Archaeplastida is marked with red boxes. In both cases, protein families were sorted by their distribution among the six eukaryotic supergroups Archaeplastida, Excavata, Hacrobia, Mycetozoa, Opisthokonta and the SAR group.

**Suppl. Fig. 3: InterProScan results for all investigated asgardarchaea as well as 14 reference archaea**. InterProScan was used to annotate the proteomes of all 12 investigated asgardarchaea and 14 reference archaea. InterProScan results were used to search for ESP candidate proteins within all analyzed proteomes and their presence indicated by a black dot. InterPro identifiers pointing towards the presence of ESPs were obtained from Zaremba et al. 2017 and in some cases revised to be more strict. For most ESPs results show a clear divide between asgardarchaea and the chosen reference archaea, while the DNA-directed RNA-Polymerases I/III and II, Cyclins or Cyclin-like proteins and the STT3 subunit of the OST complex could also be detected in the majority of the analyzed reference archaea.

**Suppl. Table 1**. EPC overview including protein family counts, protein sequence listing for each protein family and relative frequency of occurrence within protein families of all taxonomic groups present in the clustering. One protein family per line, one member each cell.

**Suppl. Table 2**. Six identified Eukaryote-Asgardarchaea unique protein families annotated via InterProScan using standard parameters and showing results from all subject databases.

**Suppl. Table 3**. Archaeal and bacterial clusterings. One protein family per line, one member each cell.

**Suppl. Table 4**. InterProScan results for all investigated Asgardarchaea, underlying Fig. 3. in addition to the list of InterPro accessions used to identify ESPs within the investigated asgardarchaeal MAGs.

**Suppl. Table 5**. Listing of all proteome assemblies used in cluster creation and BLAST hit analysis.

**Suppl. Table 6**. Species abbreviations used in Figure 2 and Figure 3.

**Suppl. Table 7**. FDR-corrected (Benjamini-Hochberg) p-values of pairwise double-sided KS tests of gene length distributions.

## Notes

### Competing Interest Statement

The authors have declared no competing interest.

