## Supplementary Material for "The asgardarchaeal-unique contribution to protein families of the eukaryotic common ancestor was 0.3%"

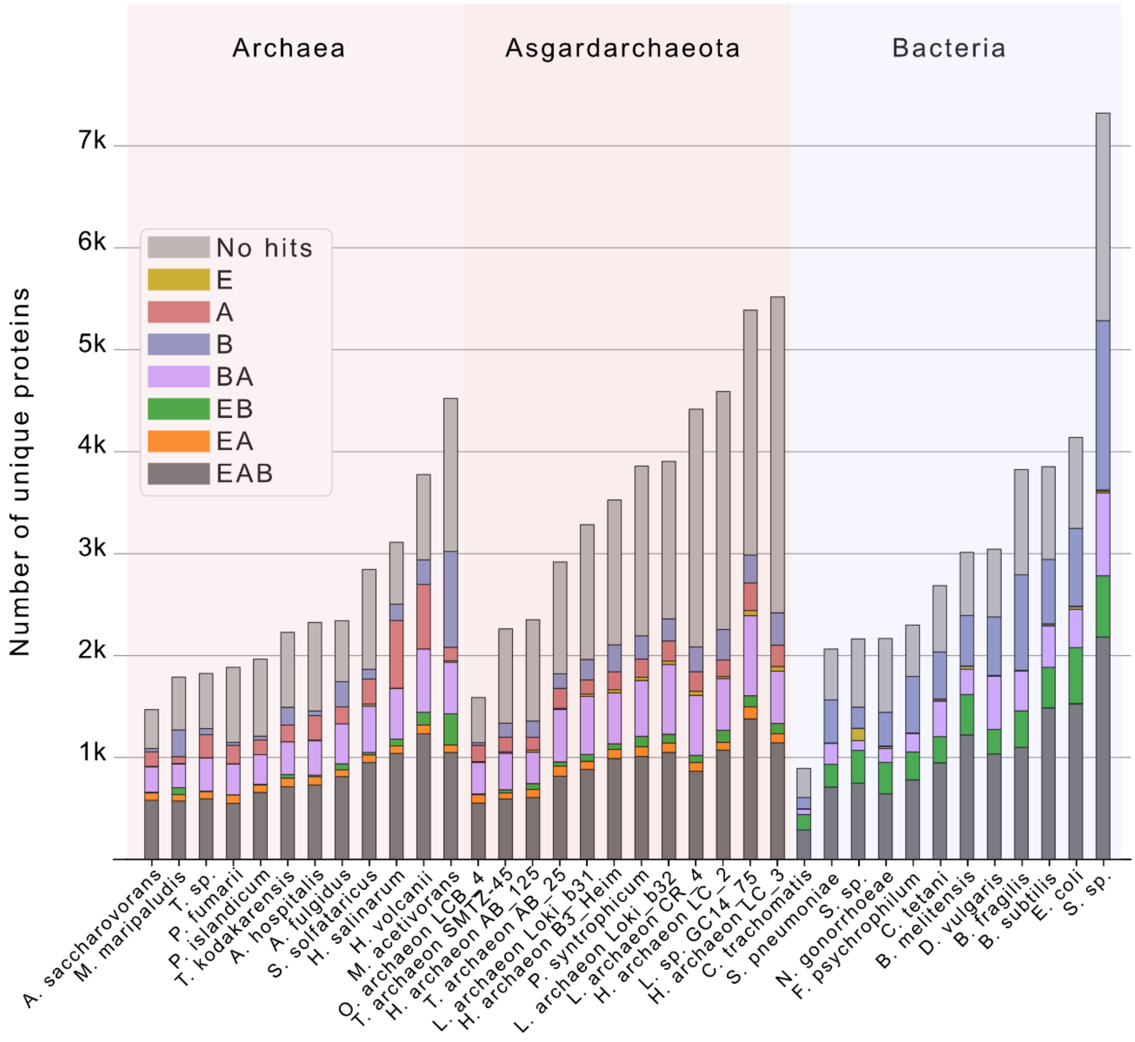


**Suppl. Fig. 1: BLAST hit analysis**. The BLAST hit analysis underlying Fig. 3a was redone, this time ignoring all hits from the same phylum. This was done in order to test for the asgardarchaeas' taxonomic rank, since the amount of protein sequences without any significant database hits, could either stem from poor sequence quality or just from the fact that asgardarchaea are inherently different from all archaea we know so far. As expected, the amount of Refseq archaeal protein sequences without significant database hits did increase but the effect was not sufficient to recover the same proportion of "unknown" protein sequences as we see in asgardarchaea.

Suppl. Fig. 2

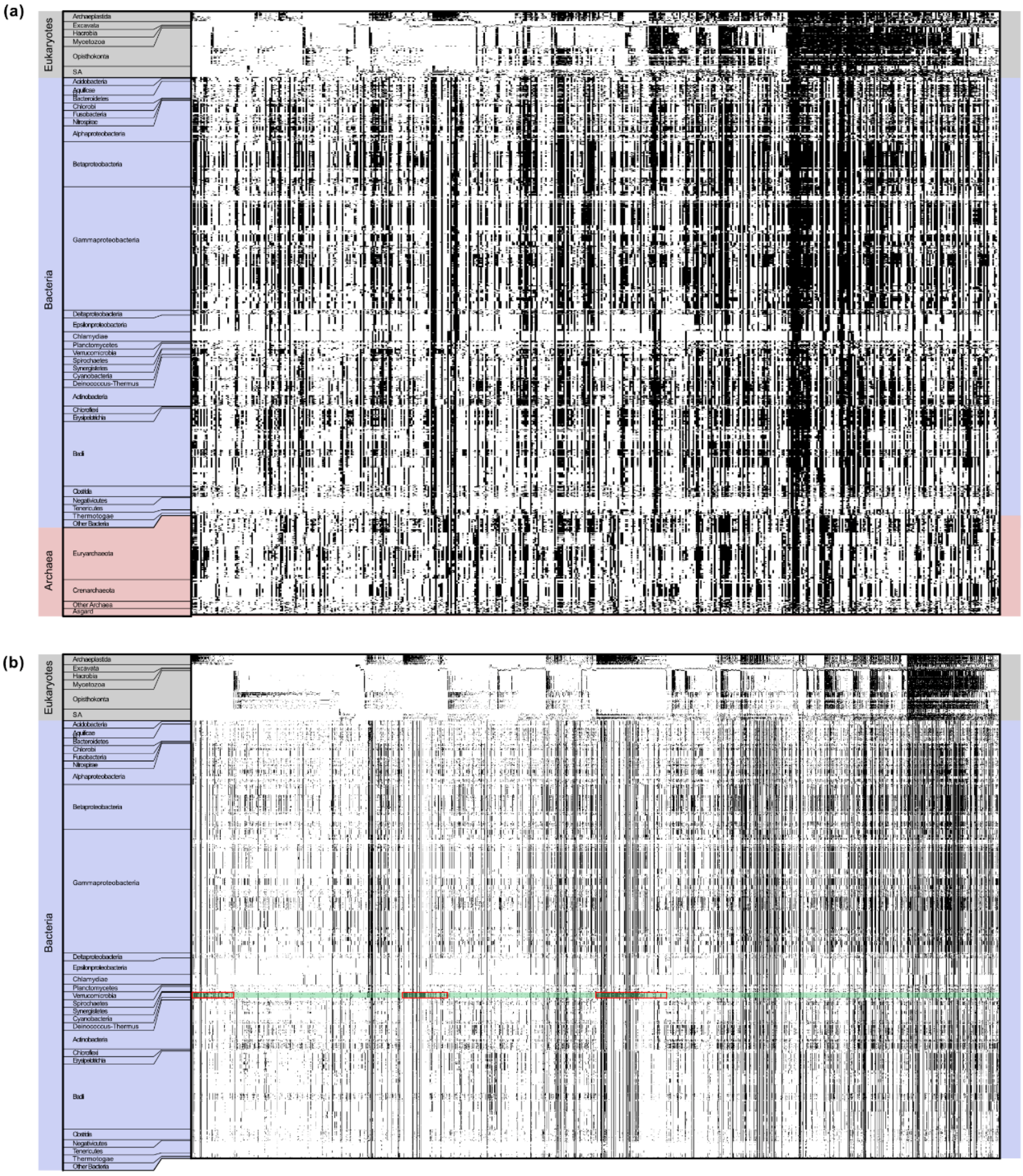


**Suppl. Fig. 2: PAPs of sets of Eukaryote-Prokaryote-Clusters. (a)** Presence-Absence pattern of all 537 EPCs containing members from eukaryotes, bacteria and archaea. **(b)** Presence-Absence pattern of all 1723 EPCs containing only proteins from eukaryotes and bacteria. The cyanobacterial contribution to the Archaeplastida is marked with red boxes. In both cases, protein families were sorted by their distribution among the 6 eukaryotic supergroups Archaeplastida, Excavata, Hacrobia, Mycetozoa, Opisthokonta and the SAR group.

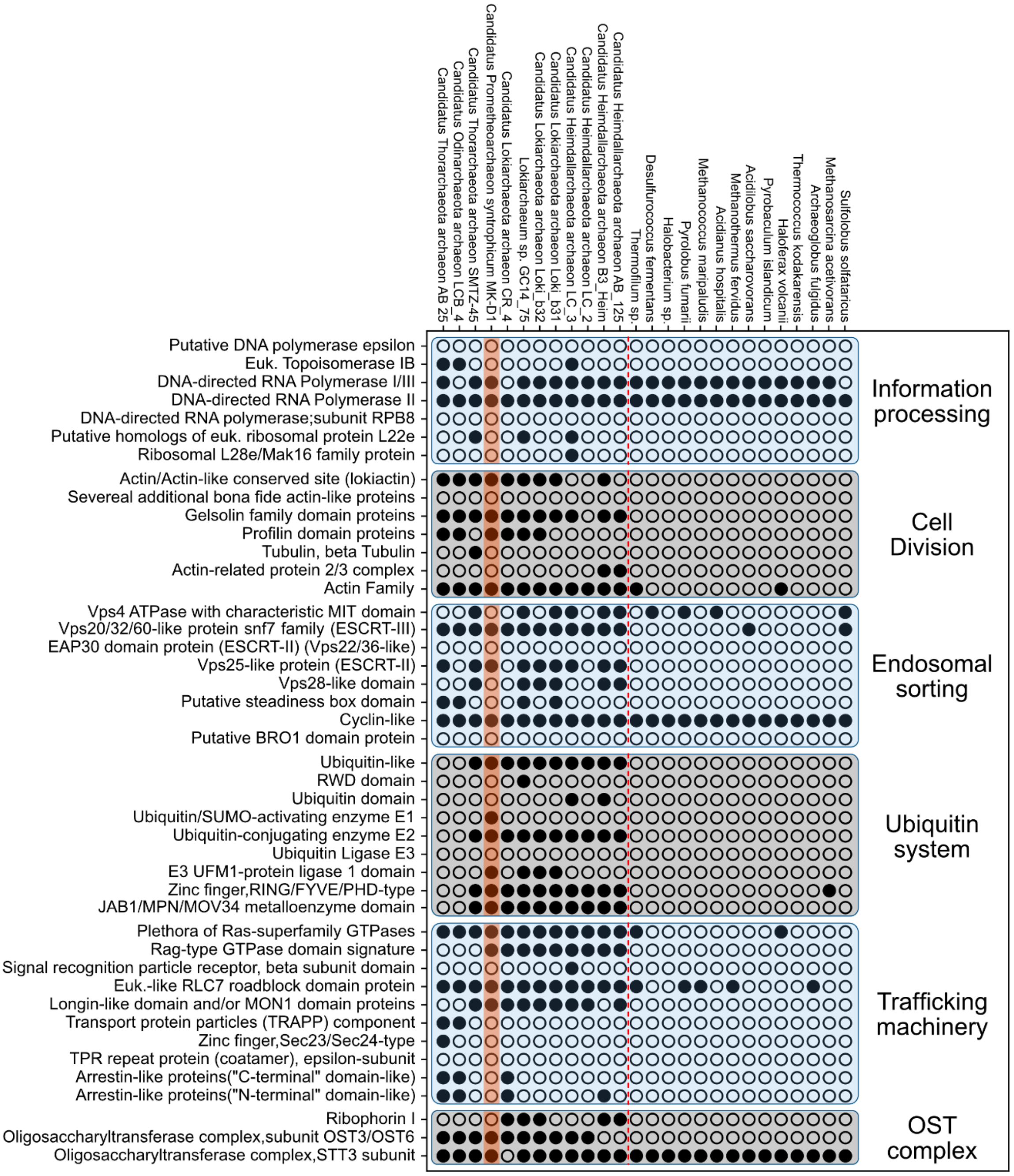


**Suppl. Fig. 3: InterProScan results for all investigated asgardarchaea as well as 14 reference archaea.** InterProScan was used to annotate the proteomes of all 12 investigated asgardarchaea and 14 reference archaea. InterProScan results were used to search for ESP candidate proteins within all analyzed proteomes and their presence indicated by a black dot. InterPro identifiers pointing towards the presence of ESPs were obtained from Zaremba et al. 2017 and in some cases revised to be more strict. For most ESPs results show a clear divide between asgardarchaea and the chosen reference archaea, while the DNA-directed RNA-Polymerases I/III and II, Cyclins or Cyclin-like proteins and the STT3 subunit of the OST complex could also be detected in the majority of the analyzed reference archaea.

**Suppl. Table 1.** EPC overview including protein family counts, protein sequence listing for each protein family and relative frequency of occurrence within protein families of all taxonomic groups present in the clustering. One protein family per line, one member each cell.

**Suppl. Table 2.** Six identified Eukaryote-Asgardarchaea unique protein families annotated via InterProScan using standard parameters and showing results from all subject databases.

**Suppl. Table 3.** Archaeal and bacterial clusterings. One protein family per line, one member each cell.

**Suppl. Table 4.** InterProScan results for all investigated Asgardarchaea, underlying Fig. 3. in addition to the list of InterPro accessions used to identify ESPs within the investigated asgardarchaeal MAGs.

**Suppl. Table 5.** Listing of all proteome assemblies used in cluster creation and BLAST hit analysis.

**Suppl. Table 6.** Species abbreviations used in Figure 2 and Figure 3.

**Suppl. Table 7.** FDR-corrected (Benjamini-Hochberg) p-values of pairwise doule sided KS tests of gene length distributions
